# Haplotype-specific chromosome painting unveils recombination patterns in the holocentric species *Rhynchospora breviuscula* H.Pfeiff

**DOI:** 10.64898/2026.06.24.733714

**Authors:** Thiago Nascimento, André Marques

## Abstract

The genus *Rhynchospora* Vahl (beak-sedges) comprises approximately 381 accepted species with a worldwide distribution, all of which possess holocentric chromosomes, where centromeric activity is distributed almost along the entire chromosome. Despite the recent advances, the mechanisms governing the dynamics of meiotic recombination in holocentric plants remain poorly understood. Here, we developed haplotype-specific oligo-FISH probes for chromosomes 1, 2, and 3 based on a haplotype-phased genome assembly of *Rhynchospora breviuscula* (*n* = 5), enabling homolog-specific chromosome painting. Each probe set was labelled with a distinct fluorophore and hybridised *in situ* to metaphase chromosomes of the reference plant and seven F_1_ individuals derived from self-crossed reference plants. This approach allowed the unambiguous discrimination of homologous haplotypes and the indirect visualisation of crossover (CO) events in recombined chromosomes. We observed that recombination events were predominantly located in terminal chromosomal regions, consistent across individuals. These results corroborate previous findings from single-cell recombination mapping and provide independent cytological validation of the recombination landscape in this species. Our study establishes haplotype-specific chromosome painting as a robust tool for high-resolution mapping of meiotic recombination in holocentric plants across generations. Furthermore, these probes provided a foundation for future investigations into inverted meiosis, a mechanism characterized by an alternative pattern of chromosome segregation in holocentric species.

## 1. Introduction

Meiotic cell division is an essential process underpinning the maintenance of biological diversity. This process is highly conserved from yeast to mammals and is characterised by a single round of DNA replication followed by two successive rounds of chromosome segregation, thereby reducing the chromosome number by half and producing haploid gametes from diploid cells[1]. Meiosis involves the programmed formation of DNA double-strand breaks (DSBs) mediated by the SPO11 protein, followed by homologous chromosome pairing and synapsis, crossover (CO) formation and recombination, and the stepwise release of sister chromatid cohesion[2–4]. Through these coordinated steps, meiosis fulfils dual and seemingly contrasting roles: preserving genome stability while simultaneously generating genetic diversity via recombination and independent assortment[5].

Variation in meiotic processes can give rise to structural rearrangements, such as deletions and duplications, thereby contributing to genome evolution; however, such variation may also result in severe defects, including infertility[6,7]. Despite the conservation of its core mechanisms, meiosis exhibits notable species-specific variation, including differences in crossover landscapes among plant species and divergence in the function of meiotic protein complexes, such as monopolin, across taxa[4,8].A deeper understanding of meiosis has enabled important agricultural applications, including the manipulation of recombination for crop improvement and the development of technologies such as *MiMe* for engineering apomeiosis, while also providing key insights into evolutionary processes[4,9,10].

In holocentric organisms, which possess kinetochores distributed along the length of the chromosome, two distinct meiotic patterns have been described. A canonical-like meiosis, similar to monocentric organisms, in which homologous chromosomes segregate during meiosis I, followed by the separation of sister chromatids in meiosis II. This conventional sequence is observed, for example, in the holocentric nematode *Caenorhabditis elegans* during oocyte meiosis[11].In contrast, inverted meiosis reverses this order, with sister chromatids separating during meiosis I and homologous chromosomes segregating during meiosis II[12,13]. Inverted meiosis appears to have evolved independently in multiple lineages and has been reported in diverse holocentric taxa, including wood white butterflies (*Leptidea sinapis*)[12], sedges (*Rhynchospora pubera, R. tenuis*, and *Luzula elegans*)[14,15], true bugs (*Graphosoma italicum*)[16], and mealybugs (*Planococcus citri*)[17]. In *L. sinapis*, evidence for inverted meiosis was inferred from the identical distribution patterns of 18S rDNA loci observed in metaphase I and II[12]. In *Rhynchospora* and *Luzula*, direct cytological observations demonstrated the segregation of sister chromatids to opposite poles during anaphase I, while homologous non-sister chromatids aligned and segregated during the second meiotic division[14,15].

Cytogenetic mapping of COs along chromosomes can be achieved using several complementary approaches. One widely used method involves the immunodetection of recombination-associated proteins, such as HEI10 and MLH1, along the synaptonemal complex during late pachytene[18]. In nascent allopolyploids, genomic *in situ* hybridisation (GISH) can be employed to detect crossovers; however, this approach is limited to identifying transitional regions between homoeologous chromosomes and is most effective when subgenomes are sufficiently diverged[19]. Alternatively, fluorescence *in situ* hybridisation (FISH) using predefined DNA markers enables the inference of CO positions based on genetic distances between markers, although it lacks fine-scale resolution[20,21]. In this context, the development of haplotype-specific chromosome painting probes represents a significant advance in CO mapping. In plants, such probes were first developed for maize (*Zea mays* L.)[22], and more recently for autotetraploid sugarcane (*Saccharum spp*.)[23] and cucumber (*Cucumis sativus* L.)[24].

Here, we extend this approach to a holocentric plant system by developing haplotype-specific oligo-FISH painting probes for *Rhynchospora breviuscula*. Using these probes, we distinguish parental haplotypes in the somatic chromosomes of F1 individuals, enabling the identification of recombinant chromosomal segments inherited from previous meiotic CO events. Thus, rather than directly visualising recombination during meiosis, our approach captures its outcomes in mitotic cells and allows the reconstruction of crossover positions along individual chromosomes. This strategy provides a cytogenetic framework for analysing recombination landscapes in species with repeat-based holocentromeres and establishes a foundation for future studies investigating the relationship between recombination and chromosome segregation in holocentric systems, including those exhibiting inverted meiosis.

## 2. Methodology

### (a) Plant material

The reference *R. brevisuscula* plants were collected from natural populations occurring in Iporanga (São Paulo state, Brazil) in 2013 and further kept cultivated in greenhouse under specific conditions (16h daylight, at 26 °C and >70% humidity). After random rounds of self crossings, the F_1_ springs were also cultivated and used.

### (b) Probes design and synthesis

In order to design haplotype-specific painting probes for *R. breviuscula*, its available phased genome was used as reference (Accession: GCA_027562975.1), following the protocol used for maize[22] with a few modifications. Probes were designed specifically for chromosomes 1, 2 and 3, since chromosomes 4 and 5 are highly homozygous, which hinders the selection of enough markers to differentiate both haplotypes. For this, the Chorus2 pipeline (https://github.com/zhangtaolab/Chorus2)[25] was applied with the following parameters “-l 45 –homology 75 –step 5”, a first round of masking was done to prevent repetitive regions and then the design of 45-bp long oligos that contemplate specific regions to differentially recognize both haplotypes. It was also applied a round of repetitiveness checking through *k*-mer analysis for both haplotypes (17 bp), excluding all the oligos with more than two *k*-mer copies.

After the design process a total of 15545 oligos were recovered for Chr1_h11, 15331 oligos for Chr1/H2, 13615 oligos for Chr2_h1, 13777 oligos for Chr2_h2, 12061 oligos for Chr3_h1 and 11847 oligos for Chr3_h2 (For the specific coordinates and sequences of each oligo, please check the data accessibility section). The oligos were then synthetized by Arbor Bioscience® company, with probes for haplotype one being labelled in green (Alexafluor 488) and for haplotype two in red (ATTO-633). Also, the immortal libraries were provided for further indirect labelling with digoxigenin and biotin through Microarray Debubbling PCR (Braz et al. 2020), later detected by secondary antibodies anti-biotin Alexafluor 488 (Invitrogen) and anti-digoxigenin rhodamine (Roche) (**Fig.1**).

### (c) Chromosomes preparation

Root tips from different individuals of *R. breviuscula* were collected and fixed in Carnoy solution (3:1 v:v, Methanol:Acetic Acid) per 2-24h at room temperature and then stored at -20 °C until the moment of use. The slides were prepared using an air-dry based technique. Briefly, the meristematic tissue was pre-treated with an enzymatic solution containing 2% cellulase-R10 Onozuka (Serva), 20% pectinase (Sigma), 2% pectolyase Y23 (Duchefa), 2% Cytohelicase (Sigma) diluted in 1× citric buffer (pH: 6.0) for 2h at 37 °C in a humid chamber. After enzymatic digestion, the excess of solution was removed and then the material was washed with distilled water. The meristem was macerated in a drop of fresh cold fixative solution with the help of needles, then added another drop and spread in the surface of the slide, followed by consecutives air pumps. After this procedure, the slides were soaked in 60% acetic acid at room temperature for 30-60 min and then left to dry in the fume hood until complete evaporation. For selecting the best slides, chromosomes were stained with 4′,6-diamidino-2-phenylindole (DAPI) in a concentration of 2 µg/mL in Vectashield antifade medium (Vector Labs) and then checked in fluorescence microscope.

### (d) Oligo-FISH

The oligo-FISH experiment was conducted following to the instructions proposed by Braz et al. (2020) with a few modifications. The hybridization mix consisted of 50% formamide, 2 × Saline Sodium Citrate solution (SSC, pH 7.0), 10% dextran sulphate, ∼ 350 ng of the probe labelled in green and ∼200 ng of the probe labelled in red, in a total volume of 10 µL per slide. The chromosomes were denatured for 5 min at 85 °C and incubated for at least 18h at 37 °C in a humid dark chamber. After that, the coverslips were removed and stringency washes were performed in 2 × and 0.1 × SSC at 42 °C (achieving a total of ∼ 76% final stringency). For the detection of indirect labelled probes, this step was followed by a wash in 1 × Tris-NaCl-Blocking buffer (TNB) and after applied 20 µL of the same solution containing 0.2 µL of rhodamine sheep anti-DIG (Roche) and 0.2 µL of Alexafluor 488 Streptavidin (Invitrogen) with posterior incubation at 37 °C for 1 h. Chromosomes were counterstained with 2 µg/mL DAPI in Vectashield antifade solution (Vector Laboratories). Images of metaphasic chromosomes were captured using a Zeiss Axio Imager Z2 with ZEN blue software (Zeiss). The best images were treated in brightness and contrast using Adobe Photoshop (2026).

## 3. Results

### (a) Identification and synthesis of haplotype-specific probes

To enable haplotype-resolved cytogenetic analyses in *Rhynchospora breviuscula*, we established chromosome-specific oligo-FISH probe sets based on the haplotype-phased reference genome assembly (∼420 Mb; Castellani & Zhang et al. 2024). Both genomes (haplotype 1 and 2) were organised into five pseudochromosomes; however, robust haplotype-specific probe design was only achievable for chromosomes 1, 2, and 3 of each haplotype. In contrast, chromosomes 4 and 5 contain extensive homozygous regions, which limited the identification of sequences uniquely assignable to each haplotype and precluded the generation of specific probe sets. For the three target chromosomes, we obtained dense sets of haplotype-specific 45-bp oligonucleotides, enabling clear discrimination between homologous chromosomal segments. These probe sets provided the basis for subsequent cytogenetic analyses of haplotype structure and recombination patterns. In total, 82176 oligos were generated, being 15545 oligos for chromosome 1/ haplotype 1 (Chr1_h1; ∼170 oligos/Mb) and 15331 for chromosome 1/ haplotype 2 (Chr1_h2; ∼172 oligos/Mb), 13615 for chromosome 2/ haplotype 1 (Chr2_h1; ∼192 oligos/Mb) and 13777 for chromosome 2/ haplotype 2 (Chr2_h2; ∼191 oligos/Mb), 12061 for chromosome 3/ haplotype 1 (Chr3_h1; ∼173 oligos/Mb) and 11847 for chromosome 3/ haplotype 2 (Chr3_h2; ∼169 oligos/Mb). After complete selection, the oligos were then synthetized in large scale and labelled (See Methods section).

### (b) Validation through oligo-FISH chromosome painting

The different pools of oligos for each haplotype were labelled with different fluorophores (haplotype 1 in green and haplotype 2 in red) and then hybridised to the metaphase chromosomes of *R. breviuscula* reference plant through oligo-FISH painting technique. Each pool of probes generated clear signals (**Fig. 1**), allowing the precise identification of both haplotypes for chromosome 1 (**Fig. 1A-C**), chromosome 2 (**Fig. 1D-F**) and chromosome 3 (**Fig. 1G-I**). However, for chromosome 2, a region of haplotype 1 was highly homozygous, causing a cross hybridization of haplotype 2 probes to this region, generation a red signal in Chr2_h1 (**Fig. 1D-F**). This region was not identified previously by the selection step with the Chorus2.

**Figure 1.**
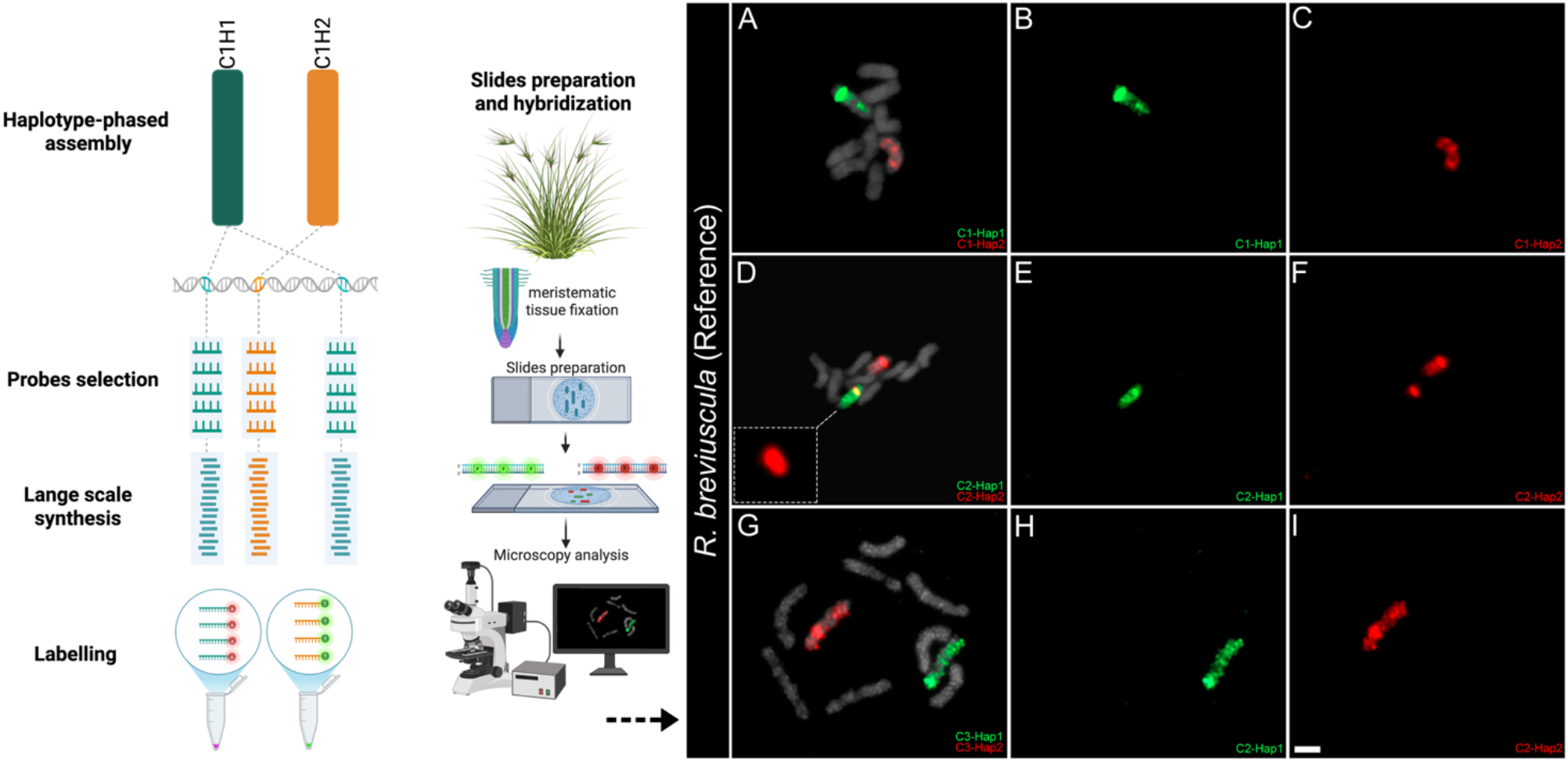
Development oh haplotype-specific probes for chromosomes 1, 2 and 3 (Chr1- 3) of *R. breviuscula*. The scheme on the left represents the main steps of the selection and construction of the oligo probes plus further slides preparation and hybridisation. On the right, oligo-FISH painting results are shown for the *R. breviuscula* reference plant, with probes for haplotype 1 (H1) of each chromosome marked in green (**B, E** and **H**), while probes for haplotype 2 (H2) of each chromosome are labelled in red (**C, F** and **I**), overlay of the probes with the chromosomes counterstained in DAPI (in grey) are also provided (**A, D** and **G**). Insert in **D** indicates the cross hybridization of probes for haplotype 2 in haplotype 1. Scale bar in **I** corresponds to 2 µm. Scheme on the left was designed using BioRender®.

### (c) Revealing CO spots through chromosome painting

Using this set of haplotype-specific probes, we investigated the distribution of crossovers across different F1 plants. The reference plant was self-pollinated to generate several F1 individuals (F1.1–F1.6), in addition to a distinct accession (*R. breviuscula* IPO). Chromosome painting experiments revealed multiple recombination hotspots resulting from exchanges between homologous chromosomes (**Fig. 2**). For chromosome 1, recombination occurred predominantly in subtelomeric regions. In the F1-1, F1-2 and F1-4 plants, exchanges were largely reciprocal between homologous chromosomes. In contrast, in F1-3 and F1-5 the exchange occurred only in one homolog: haplotype 2 (red) contained a segment derived from haplotype 1 (green). In F1-6, the opposite configuration was observed, with a segment of haplotype 2 incorporated into haplotype 1. Chromosome 2 also exhibited several chromosomal exchanges. In the F1-1 and F1-4 plants, recombination occurred at different positions along the chromosome. In F1-2, haplotype 1 was almost entirely replaced by haplotype 2, with only a small terminal segment of haplotype 1 remaining. A similar pattern was observed in F1-6, although in this case haplotype 2 was nearly completely replaced by haplotype 1. In F1-3 and F1-5, a comparable configuration was detected, with homolog 1 containing a small terminal segment derived from haplotype 2.

**Figure 2.**
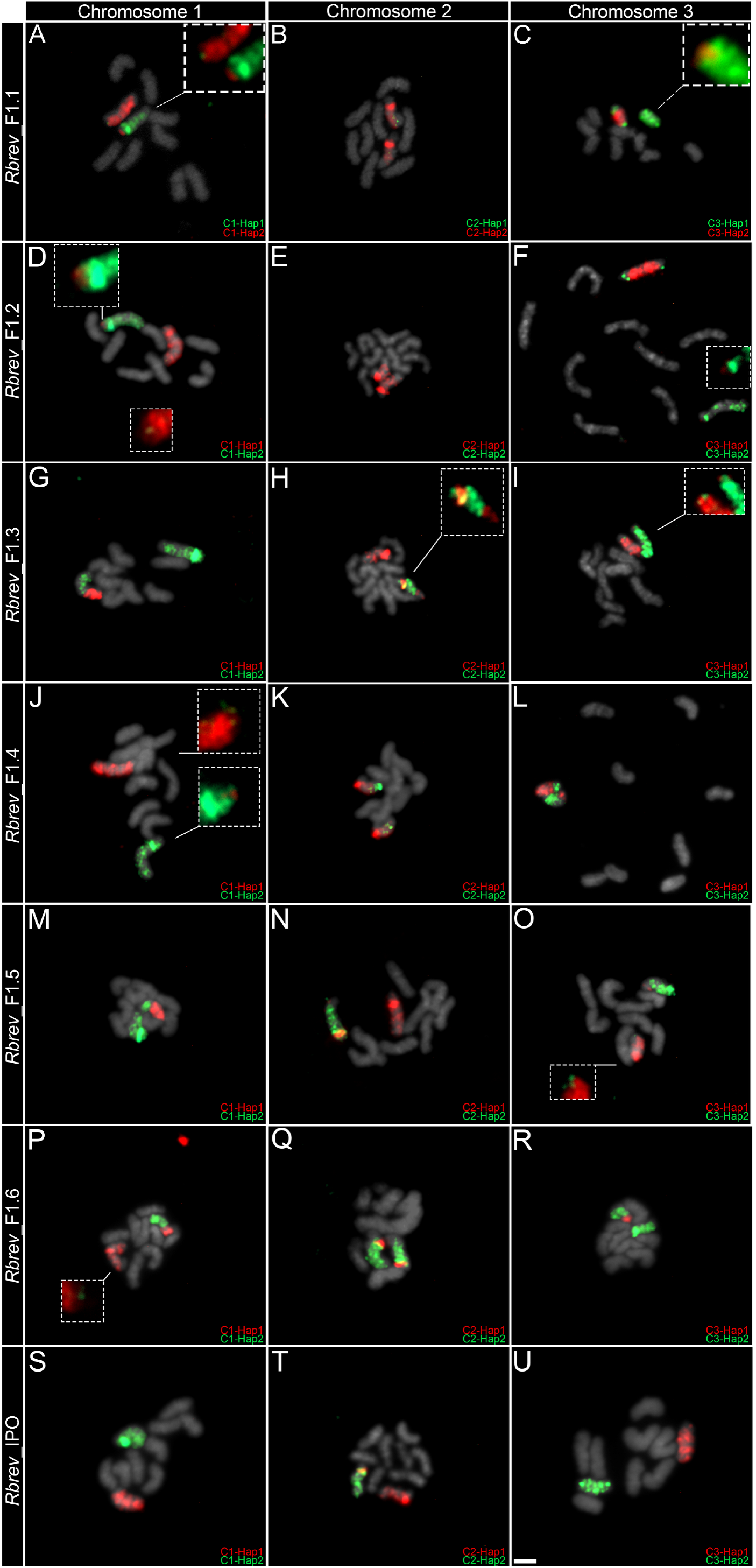
Oligo-FISH chromosome painting with haplotype-specific probes for chromosomes 1 (Chr1_h1-h2), 2 (Chr2_h1-h2) and 3(Chr3_h1-h2) in different F1 plants of R. *breviuscula*. Probes for haplotype 1 of each chromosome are represented in green, while for haplotype 2 are represented in green. The different F_1_ plants are treated here as: F1-1 (**A**-**C**), F1-2 (**D**-**F**), F1-3 (**G**-**I**), F1-4 (**J**-**L**), F1-5 (**M**-**O**), F1-6 (**P**-**R**). The *R. breviuscula* “Iporanga” accession is treated as IPO (**S**-**U**). Bar in **U** corresponds to 2 µm.

Analysis of chromosome 3 revealed similar patterns in the F1-1 and F1-2 plants. In these individuals, homolog 1 displayed a small signal corresponding to haplotype 2 at one terminal region, whereas homolog 2 contained haplotype 1 signals at both termini. In F1- 3, F1-4 and F1-5, exchanges were reciprocal, with both homologs exhibiting exchanged signals in one terminal region, although F1-4 displayed a slightly larger exchanged segment. In F1-6, most of homolog 2 was replaced by haplotype 1 sequences, with only a single terminal haplotype 2 signal remaining.

In contrast, the IPO accession showed no detectable differences compared with the reference plant, suggesting that these two individuals share nearly identical genomic haplotypes. Overall, recombination hotspots in *R. breviuscula* were consistently located in the terminal regions of the chromosomes (**Fig. 3 and 4**).

**Figure 3.**
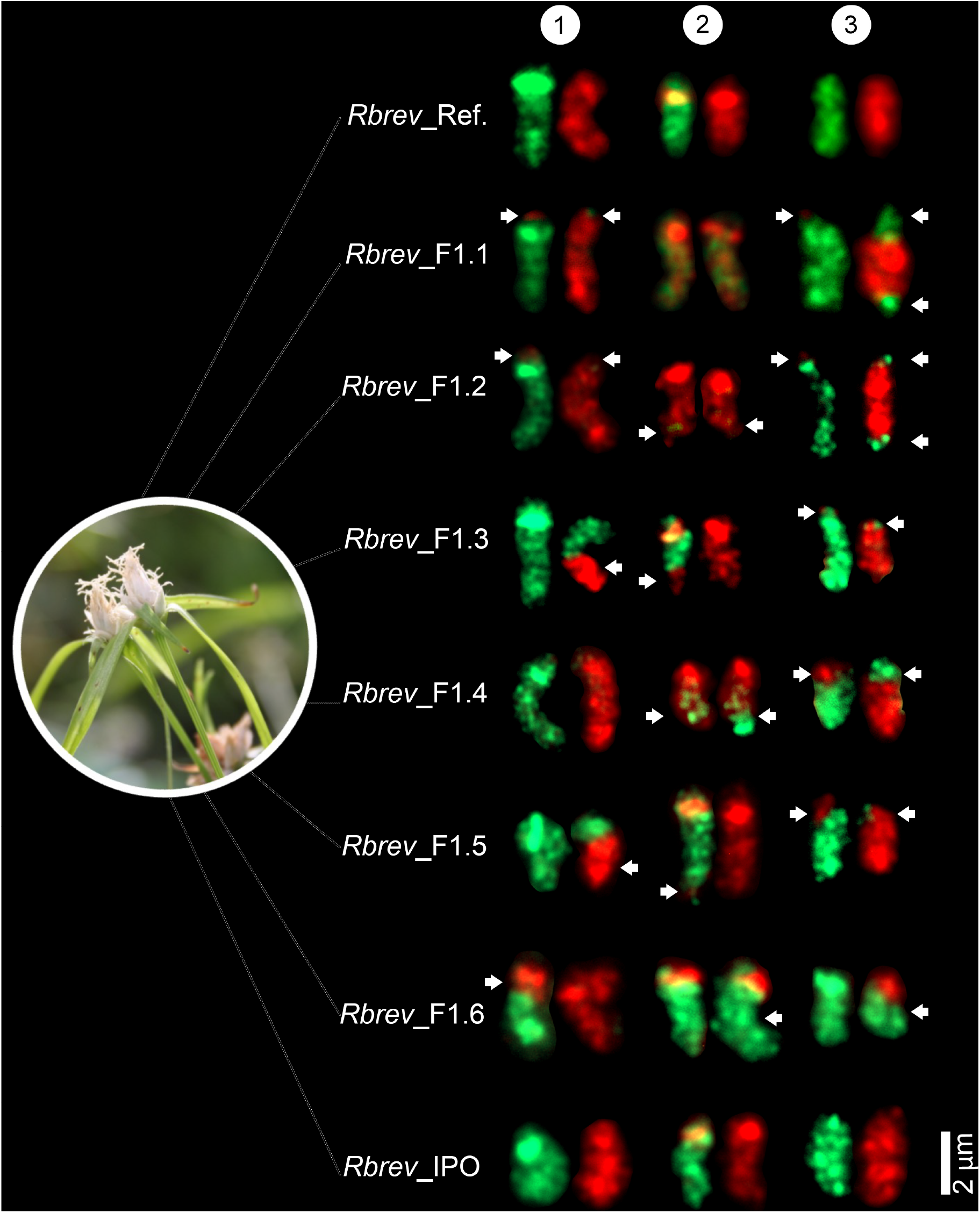
Karyogram detailing all the chromosomal exchange regions between haplotypes 1 and 2. Haplotype 1 probes are shown in green, whereas haplotype 2 are shown in red. Arrows indicate the hotspots of recombination and small signals of exchange.

**Figure 4.**
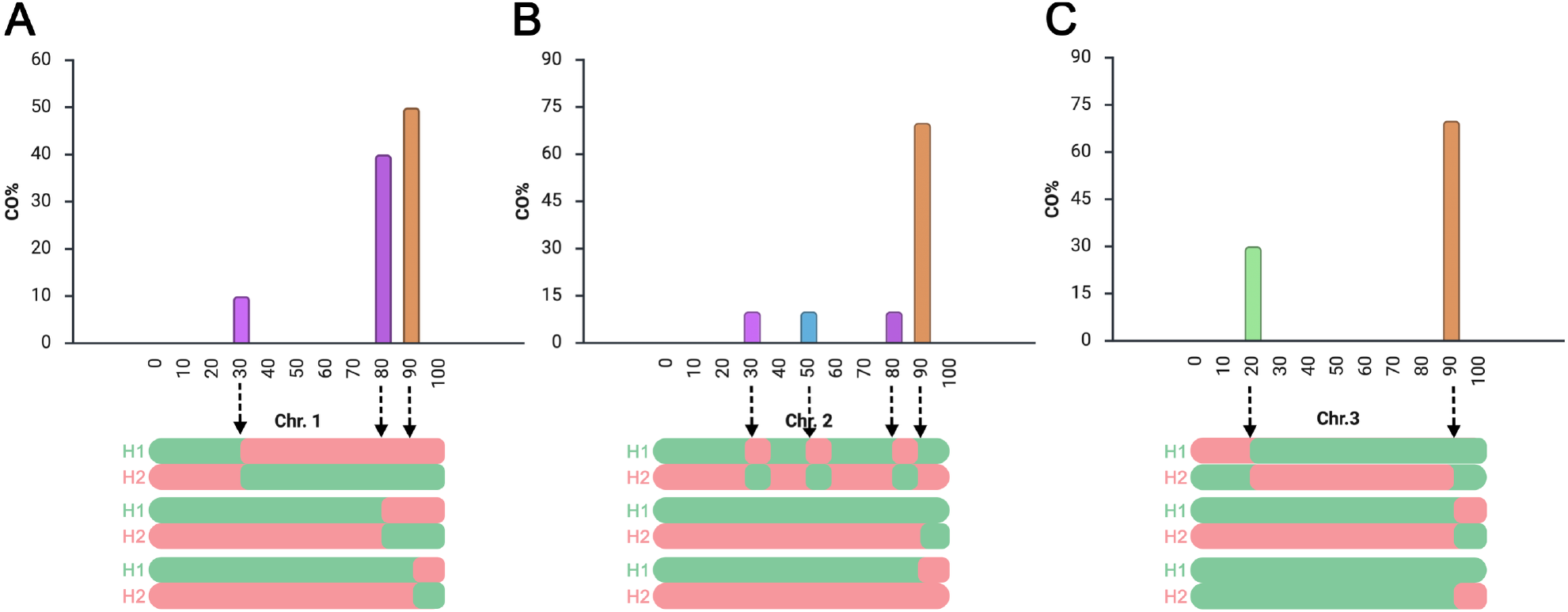
Distribution and frequency of meiotic crossovers (COs) on chromosomes 1 (**A**), 2 (**B**), and 3 (**C**) in F_1_ individuals of *R. breviuscula*. Chromosomes were normalized and divided into 100 equal intervals (0–100) along their length, represented on the x-axis. Meiotic COs identified in **Figure 3** were mapped to their corresponding intervals. The y-axis indicates the frequency of COs within each interval, calculated as the frequency of COs per interval divided by the total number of chromosomes analysed. Schematic representations of the observed exchange configurations between haplotypes are shown below each frequency plot (haplotype 1 in green; haplotype 2 in red)

## 4. Discussion

The number and distribution of CO events along plant chromosomes are tightly regulated. In most species, one to three COs occur per chromosome and are preferentially located in distal regions. In monocentric chromosomes, this pattern is commonly associated with the presence of pericentromeric heterochromatin, which is enriched in tandem repeats and transposable elements and is generally refractory to crossover formation, thereby contributing to reduced recombination in pericentromeric regions[26– 28]. In contrast, holocentric chromosomes, lack a localized centromere, raising the expectation that recombination suppression might be more uniformly distributed along their length. However, the mechanisms governing recombination patterning in holocentric plants remain poorly understood.

In the genus *Rhynchospora*, holocentromeres of most species are associated with the satellite DNA *Tyba*, which is dispersed along chromosomes[13,29–31], a feature that could potentially lead to widespread suppression of recombination. Surprisingly, recent recombination maps based on single-cell approaches in *R. breviuscula* revealed a pattern similar to those observed in monocentric species, with one or two COs per chromosome predominantly located in terminal regions[32]. Our haplotype-specific chromosome painting results independently corroborate these findings at the cytological level and further demonstrate that this distal bias is consistent across multiple individuals.

The persistence of terminally biased recombination in a holocentric context suggests that factors other than centromere localization play a dominant role in shaping CO landscapes. Possible explanations include large-scale chromatin organisation, such as differences in chromatin accessibility or epigenetic state along chromosome arms, as well as constraints imposed by the meiotic chromosome axis and crossover interference[33]. Disentangling these possibilities will require the integration of cytological, genomic, and epigenetic datasets, but our results provide a critical framework for such analyses.

Comparative analyses across holocentric taxa suggest that the absence of a localized centromere does not preclude the emergence of conserved strategies to ensure accurate meiotic chromosome behaviour. Studies in the lepidopteran model *Bombyx mori* using oligo-FISH painting probes have shown that homologous chromosomes preferentially initiate pairing at telomeric regions, implying that chromosome ends may function as key spatial cues for homolog recognition and synapsis in holocentric systems[34]. Broader comparative work across Lepidoptera indicates that this telomere-driven pairing architecture is evolutionarily conserved and associated with the formation of elongated rod-like bivalents during meiosis I[35]. Notably, despite the diffuse kinetochore organization typical of holocentric chromosomes, these species appear to establish transiently localized spindle interactions, effectively generating a functional “centromere-like” domain during segregation. Similar solutions may operate in holocentric plants such as *Rhynchospora* and *Luzula*, where specialized meiotic configurations, including inverted meiosis, suggest that diverse lineages have independently evolved mechanisms that spatially reorganize chromosome architecture to preserve faithful segregation despite fundamentally different centromere organization[14,15].

The availability of haplotype-specific probes for chromosome painting in plants remains limited to a small number of species, largely due to the requirement for haplotype-phased genome assemblies to enable the accurate selection of haplotype-specific oligonucleotides. However, this scenario has been progressively changing with advances in sequencing technologies and bioinformatic tools. The first haplotype-specific oligo-FISH experiment in plants was conducted in maize (*Zea mays*), where probes were designed to distinguish the B73 and Mo17 haplotypes of chromosome 10 and to analyse CO distribution across F_1_ and F_2_ hybrids, besides recombinant inbred lines [22]. In autotetraploid sugarcane (*Saccharum* spp.), this approach enabled the discrimination of all four subgenomes and revealed the absence of major structural rearrangements among homologous chromosomes, thereby establishing a powerful tool for investigating haplotype-specific chromosome structure in polyploid species[23]. More recently, a similar system was developed for cucumber (*Cucumis sativus*), revealing preferential crossover hotspots along chromosome arms and differences in recombination rates between cultivated and more ancestral accessions. This study also enhanced the methodology through the use of secondary oligonucleotides to increase signal intensity, an improvement that is particularly valuable when only a limited number of informative oligos can be selected, such as in species with small or highly repetitive genomes[24]. By extending this methodology to a holocentric plant species, our study broadens its applicability and establishes haplotype-specific cytogenetics as a versatile tool for investigating chromosome behaviour in non-canonical systems.

Inverted meiosis is an important feature associated with holocentric chromosome organization. In plants, it has so far been reported only in species of *Rhynchospora* (e.g., *Rhynchospora pubera* and *Rhynchospora tenuis*)[14] and in *Luzula elegans*[15]. In *Rhynchospora*, both chiasmatic and achiasmatic forms of inverted meiosis have been described from cytological analyses, revealing that sister chromatids segregate during meiosis I, followed by the alignment and segregation of non-sister chromatids in meiosis II. In *L. elegans*, homologous non-sister chromatids remain connected by satellite DNA from metaphase I to metaphase II, facilitating this inverted order of segregation. The future application of haplotype-specific chromosome painting probes has the potential to address several unresolved aspects of meiosis in holocentric species. In particular, such approach may enable the tracking of individual chromatids across meiotic divisions, providing a framework to test whether sister chromatids reassociate with homologous partners after the first division and how they reorient on the spindle between divisions. More broadly, these tools could help clarify how recombination outcomes are linked to chromatid behaviour during the two meiotic divisions, including their contribution to segregation stability during inverted meiosis.

## 5. Conclusion

In summary, we developed the first haplotype-specific oligo-FISH chromosome painting probes for the holocentric plant *R. breviuscula*, enabling high-resolution discrimination of homologous chromosomes and the visualisation of exchanged segments between haplotypes caused by previous CO events. Using this approach, we demonstrate that recombination is consistently biased towards terminal chromosomal regions, despite the diffuse centromeric organisation characteristics of holocentric chromosomes. These findings provide independent cytological validation of previously reported recombination landscapes and highlight the conservation of distal CO patterning across diverse chromosomal architectures. Collectively, our study provides both a methodological advance and a conceptual framework for exploring meiotic processes and genome evolution in holocentric.

## Ethics

This work did not require ethical approval from a human subject or animal welfare committee.

## Data Accessibility

*R. breviuscula* haplotype-specific oligos for Chromosome 1/ Haplotype 1

*R. breviuscula* haplotype-specific oligos for Chromosome 1/ Haplotype 2

*R. breviuscula* haplotype-specific oligos for Chromosome 2/ Haplotype 1

*R. breviuscula* haplotype-specific oligos for Chromosome 2/ Haplotype 2

*R. breviuscula* haplotype-specific oligos for Chromosome 3/ Haplotype 1

*R. breviuscula* haplotype-specific oligos for Chromosome 3/ Haplotype 2

## Declaration of AI use

We have not used AI-assisted technologies in creating this article

## Author’s contribution

**TN**: Performed the experiments and wrote the paper. **AM**: Conceptualization provided resources and wrote the paper. All authors approved the final version of the manuscript.

## Conflict of interest

We declare no conflict of interests.

## Acknowledgements

This study was funded by the Max Planck Society (core funding to A.M.), and the European Union (European Research Council Starting Grant, HoloRECOMB, grant no. 101114879 to A.M.). The DFG also funded this work under Germany’s Excellence Strategy—EXC 493 2048/1–390686111 (to A.M.).

## Notes

### Competing Interest Statement

The authors have declared no competing interest.

## References

1. Ur SN, Corbett KD. 2021 Architecture and Dynamics of Meiotic Chromosomes. Annu. Rev. Genet. 55, 497–526. (doi:10.1146/annurev-genet-071719-020235)

2. Roeder GS. 1997 Meiotic chromosomes: it takes two to tango. Genes Dev. 11, 2600–2621. (doi:10.1101/gad.11.20.2600)

3. Marston AL, Amon A. 2004 Meiosis: cell-cycle controls shuffle and deal. Nat. Rev. Mol. Cell Biol. 5, 983–997. (doi:10.1038/nrm1526)

4. Kuo P, Da Ines O, Lambing C. 2021 Rewiring Meiosis for Crop Improvement. Front. Plant Sci. 12, 708948. (doi:10.3389/fpls.2021.708948)

5. Hultén M, Baker H, Tankimanova M. 2005 Meiosis and meiotic errors. In Encyclopedia of Genetics, Genomics, Proteomics and Bioinformatics (eds LB Jorde, PFR Little, MJ Dunn, S Subramaniam), Wiley. (doi:10.1002/047001153X.g102206)

6. Cai X, Xu S. 2007 Meiosis-Driven Genome Variation in Plants. Curr. Genomics 8, 151–161. (doi:10.2174/138920207780833847)

7. Börner GV, Cha RS. 2015 Analysis of Recombination and Chromosome Structure during Yeast Meiosis. Cold Spring Harb. Protoc. 2015, pdb.top077636. (doi:10.1101/pdb.top077636)

8. Ohkura H. 2015 Meiosis: An Overview of Key Differences from Mitosis. Cold Spring Harb. Perspect. Biol. 7, a015859. (doi:10.1101/cshperspect.a015859)

9. d’Erfurth I, Jolivet S, Froger N, Catrice O, Novatchkova M, Mercier R. 2009 Turning Meiosis into Mitosis. PLoS Biol. 7, e1000124. (doi:10.1371/journal.pbio.1000124)

10. Wang Y et al. 2024 Harnessing clonal gametes in hybrid crops to engineer polyploid genomes. Nat. Genet. 56, 1075–1079. (doi:10.1038/s41588-024-01750-6)

11. Schvarzstein M, Wignall SM, Villeneuve AM. 2010 Coordinating cohesion, co-orientation, and congression during meiosis: lessons from holocentric chromosomes. Genes Dev. 24, 219–228. (doi:10.1101/gad.1863610)

12. Lukhtanov VA, Dincă V, Friberg M, Šíchová J, Olofsson M, Vila R, Marec F Wiklund. 2018 Versatility of multivalent orientation, inverted meiosis, and rescued fitness in holocentric chromosomal hybrids. Proc. Natl. Acad. Sci. 115. (doi:10.1073/pnas.1802610115)

13. Hofstatter PG et al. 2022 Repeat-based holocentromeres influence genome architecture and karyotype evolution. Cell 185, 3153-3168.e18. (doi:10.1016/j.cell.2022.06.045)

14. Cabral G, Marques A, Schubert V, Pedrosa-Harand A, Schlögelhofer P. 2014 Chiasmatic and achiasmatic inverted meiosis of plants with holocentric chromosomes. Nat. Commun. 5, 5070. (doi:10.1038/ncomms6070)

15. Heckmann S, Jankowska M, Schubert V, Kumke K, Ma W, Houben A. 2014 Alternative meiotic chromatid segregation in the holocentric plant Luzula elegans. Nat. Commun. 5, 4979. (doi:10.1038/ncomms5979)

16. Viera A, Page J, Rufas JS. 2009 Inverted Meiosis: The True Bugs as a Model to Study. In Genome Dynamics (eds R Benavente, J-N Volff), pp. 137–156. Basel: KARGER. (doi:10.1159/000166639)

17. Bongiorni S, Fiorenzo P, Pippoletti D, Prantera G. 2004 Inverted meiosis and meiotic drive in mealybugs. Chromosoma 112, 331–341. (doi:10.1007/s00412-004-0278-4)

18. Chelysheva L et al. 2012 The Arabidopsis HEI10 Is a New ZMM Protein Related to Zip3. PLoS Genet. 8, e1002799. (doi:10.1371/journal.pgen.1002799)

19. Zhang W, Zhu X, Zhang M, Chao S, Xu S, Cai X. 2018 Meiotic homoeologous recombination-based mapping of wheat chromosome 2B and its homoeologues in Aegilops speltoides and Thinopyrum elongatum. Theor. Appl. Genet. 131, 2381–2395. (doi:10.1007/s00122-018-3160-0)

20. Cheng Z, Presting GG, Buell CR, Wing RA, Jiang J. 2001 High-Resolution Pachytene Chromosome Mapping of Bacterial Artificial Chromosomes Anchored by Genetic Markers Reveals the Centromere Location and the Distribution of Genetic Recombination Along Chromosome 10 of Rice. Genetics 157, 1749–1757. (doi:10.1093/genetics/157.4.1749)

21. Tang X et al. 2008 Cross-Species Bacterial Artificial Chromosome–Fluorescence in Situ Hybridization Painting of the Tomato and Potato Chromosome 6 Reveals Undescribed Chromosomal Rearrangements. Genetics 180, 1319–1328. (doi:10.1534/genetics.108.093211)

22. Martins L do V et al. 2019 Meiotic crossovers characterized by haplotype-specific chromosome painting in maize. Nat. Commun. 10, 4604. (doi:10.1038/s41467-019-12646-z)

23. Meng Z, Shi S, Shen H, Xie Q, Li H. 2023 Haplotype-specific chromosome painting provides insights into the chromosomal characteristics in self-duplicating autotetraploid sugarcane. Ind. Crops Prod. 202, 117085. (doi:10.1016/j.indcrop.2023.117085)

24. Zhao Q, Xiong Z, Cheng C, Wang Y, Feng X, Yu X, Lou Q, Chen J. 2025 Meiotic crossovers revealed by differential visualization of homologous chromosomes using enhanced haplotype oligo-painting in cucumber. Plant Biotechnol. J. 23, 887–899. (doi:10.1111/pbi.14546)

25. Zhang T, Liu G, Zhao H, Braz GT, Jiang J. 2021 Chorus2: design of genome-scale oligonucleotide-based probes for fluorescence in situ hybridization. Plant Biotechnol. J. 19, 1967–1978. (doi:10.1111/pbi.13610)

26. Zhang X, Hu Y, Huang K, Wright SI, Rieseberg LH. 2026 Recombination suppression in plant adaptation and speciation. New Phytol., nph.71026. (doi:10.1111/nph.71026)

27. Termolino P, Falque M, Aiese Cigliano R, Cremona G, Paparo R, Ederveen A, Martin OC, Consiglio FM, Conicella C. 2019 Recombination suppression in heterozygotes for a pericentric inversion induces the interchromosomal effect on crossovers in Arabidopsis. Plant J. 100, 1163–1175. (doi:10.1111/tpj.14505)

28. Fernandes JB, Wlodzimierz P, Henderson IR. 2019 Meiotic recombination within plant centromeres. Curr. Opin. Plant Biol. 48, 26–35. (doi:10.1016/j.pbi.2019.02.008)

29. Marques A et al. 2015 Holocentromeres in Rhynchospora are associated with genome-wide centromere-specific repeat arrays interspersed among euchromatin. Proc. Natl. Acad. Sci. 112, 13633–13638. (doi:10.1073/pnas.1512255112)

30. Costa L, Marques A, Buddenhagen CE, Pedrosa-Harand A, Souza G. 2023 Investigating the diversification of holocentromeric satellite DNA Tyba in Rhynchospora (Cyperaceae). Ann. Bot. 131, 813–825. (doi:10.1093/aob/mcad036)

31. Włodzimierz P et al. 2026 Pangenome analysis reveals the evolutionary dynamics of repeat-based holocentromeres. (doi:10.64898/2026.01.17.700053)

32. Castellani M et al. 2024 Meiotic recombination dynamics in plants with repeat-based holocentromeres shed light on the primary drivers of crossover patterning. Nat. Plants 10, 423–438. (doi:10.1038/s41477-024-01625-y)

33. Zhang M et al. 2026 A holocentric pangenome links karyotype evolution to meiotic recombination. (doi:10.64898/2026.01.17.700048)

34. Rosin LF, Gil J, Drinnenberg IA, Lei EP. 2021 Oligopaint DNA FISH reveals telomere-based meiotic pairing dynamics in the silkworm, Bombyx mori. PLOS Genet. 17, e1009700. (doi:10.1371/journal.pgen.1009700)

35. Hockens C, Lorenzi H, Wang TT, Lei EP, Rosin LF. 2024 Chromosome segregation during spermatogenesis occurs through a unique center-kinetic mechanism in holocentric moth species. PLOS Genet. 20, e1011329. (doi:10.1371/journal.pgen.1011329)

